# Resistance towards and biotransformation of *Pseudomonas*-produced secondary metabolites during community invasion

**DOI:** 10.1101/2023.06.20.545698

**Authors:** Morten L. Hansen, Zsófia Dénes, Scott A. Jarmusch, Mario Wibowo, Carlos N. Lozano-Andrade, Ákos T. Kovács, Mikael L. Strube, Aaron J. C. Andersen, Lars Jelsbak

## Abstract

The role of antagonistic secondary metabolites produced by *Pseudomonas protegens* in suppression of soil-borne phytopathogens has been clearly documented. However, their contribution to the ability of *P. protegens* to establish in soil and rhizosphere microbiomes remains ambiguous. Here, we use a four-species synthetic community to determine how antibiotic production contributes to *P. protegens* community invasion and identify community traits that alter the abundance of key *P. protegens* antimicrobial metabolites (DAPG, pyoluteorin and orfamide A). Surprisingly, mutants deficient in antimicrobial production caused similar perturbations in community composition compared to invasion by wildtype *P. protegens*. Intriguingly, while pyoluteorin and orfamide A are secreted at levels toxic to individual bacterial strains, community-level resistance circumvents toxicity. Here, we identify the underlying mechanism by which the cyclic lipopeptide, orfamide A, is inactivated and degraded by *Rhodococcus globerulus* D757 and *Stenotrophomonas indicatrix* D763. Altogether, the demonstration that the synthetic community constrains *P. protegens* invasion by detoxifying its antibiotics may provide a mechanistic explanation to inconsistencies in biocontrol effectiveness *in situ*.

## Introduction

The use of bacteria and their antimicrobial metabolites has emerged as a potential alternative to synthetic agrochemicals in control of phytopathogens. In *Pseudomonas protegens*, a model organism for biocontrol, secondary metabolites such as 2,4-diacetylphloroglucinol (DAPG), pyoluteorin, and orfamide A [1–5] have been shown to play essential roles in suppression of plant pathogens. For example, DAPG suppress the fungal take-all disease in cereal plants [6], pyoluteorin is involved in the suppression of *Pythium* damping-off of cress [4], and orfamide A can reduce bacterial wilt disease in tomato [7]. However, as bacterial biocontrol is often associated with varying efficiencies across fields [8] and plants [4], it remains an important challenge to identify and mitigate the processes responsible for these variations.

Although biocontrol is a complex phenomenon that relies on multiple processes for its effectiveness [9], central processes involves interactions between an invading biocontrol strain and its secreted antibiotics, and the resident microbial community. For example, invasion and competition with existing microbial populations in the soil and rhizosphere is a prerequisite for effective biocontrol. Although recent results have shown that specific *Pseudomonas*-produced secondary metabolites (i.e. DAPG and pyoverdine) [10] are indeed required for efficient invasion of the rhizosphere of *Arabidopsis* roots and alters the structure of the resident, synthetic microbial community (SynCom), other studies have found microbial communities that are more resilient towards invasion. For example, earlier studies have shown that *P. protegens* inoculants, including engineered variants with enhanced production of DAPG and pyoluteorin, had no impact on the structure of the bacterial community on cucumber roots or on the proportion of bacteria that were tolerant or sensitive to DAPG and pyoluteorin [11]. Similarly, inoculation with various species of secondary metabolite producing fluorescent *Pseudomonas* may result in vastly different outcomes, ranging from temporary, spatially limited, and transient effects on the natural rhizosphere microbiomes [12–15] to significant perturbations within indigenous microbiomes [16, 17]. Although these results suggest that resilience towards invading *Pseudomonas* biocontrol strains and their antibiotics may depend on the composition and activity of the invaded microbial community, little is known about the underlying community traits or mechanisms that may operate to tolerate otherwise toxic levels of antibiotics and to constrain *Pseudomonas* invasion.

In this study we use a hydrogel-based bead system [18] as *in vitro* model system to systematically explore the contribution of the secondary metabolites, DAPG, pyoluteorin, and orfamide A, to the ability of *P. protegens* DTU9.1 to invade and establish in a four-membered SynCom of commonly isolated soil bacteria [19]. The hydrogel environment has been shown to mimic soil characteristics allowing for spatial distribution of microbes, as well as surface colonization [18], and thus enabling systematic analyses of community-level interactions affecting secondary metabolism in *P. protegens* DTU9.1 in an artificial yet soil-like environment. We showed that community traits affect SynCom susceptibility towards the toxic antibiotics, and that one of the underlying mechanisms involved is hydrolysis and subsequently degradation of the cyclic lipopeptide orfamide A by several members of the four-species SynCom. These results provide insight into how levels and activity of antibiotic metabolites from invading *Pseudomonas* strains may be shaped by interspecies interactions in microbial communities and provide a framework for studying community mechanisms that affect invasion or efficacy of biocontrol strains.

## Results

### *P. protegens* DTU9.1 invades a four-membered synthetic microbial community in a soil-mimicking environment

To explore the microbial interactions and their effects on community composition over time during invasion of *P. protegens* DTU9.1, a porous hydrogel bead system was chosen as cultivation system [18], as it has been shown to mimic soil characteristics [18, 20]. The SynCom comprised co-isolated species from a soil sample site that we previously demonstrated to also contain *P. protegens* [21]. Initially, we sought to investigate the sensitivity of each SynCom member towards the three antimicrobial *Pseudomonas*-produced metabolites, DAPG, pyoluteorin, and orfamide A. *Rhodococcus globerulus* D757 displayed sensitivity towards all three metabolites, whereas three members (*Pedobacter* sp. D749, *R. globerulus* D757 and *Chryseobacterium* sp. D764) were equally susceptible to pyoluteorin with a minimal inhibitory concentration (MIC) of 8 µg/ml (Table 1). The last SynCom member, *Stenotrophomonas indicatrix* D763, was generally more resistant towards all three antimicrobial metabolites (Table 1). Additionally, we tested the susceptibility of each SynCom member towards DAPG, pyoluteorin, and orfamide A utilizing an *in vitro* inhibition assay on agar surfaces (Figure S1). The antibiotic activity of pyoluteorin towards all members was confirmed, whereas antibiosis of *R. globerulus* D757 caused by orfamide A was not observed in this experimental setup.

**Table 1.** Minimal inhibitory concentrations of DAPG, pyoluteorin, and orfamide A against the SynCom members

|  | Minimal Inhibitory Concentration<br>(µg/ml) |  |  |
| --- | --- | --- | --- |
|  | DAPG | Pyoluteorin | Orfamide A |
| *Pedobacter* sp. D749 | 16 | 8 | > 64 |
| *Rhodococcus globerulus* D757 | 8 | 8 | 16 |
| *Stenotrophomonas indicatrix* D763 | > 64 | 16 | > 64 |
| *Chryseobacterium* sp. D764 | 64 | 8 | > 64 |

Next, it was determined whether *P. protegens* DTU9.1 could colonize and maintain itself in the artificial soil-like environment when cultivated axenically (Figure S2A). After four days of cultivation, the system reached a plateau of approximately 1 · 10^8^ CFU/ml, which was maintained at the seventh and final day of the experiment. Furthermore, knock-out mutants incapable of producing DAPG (Δ*phlACB*), pyoluteorin (Δ*pltA*), or orfamide A (Δ*ofaA*) were constructed and tested for their growth within the hydrogel bead system. The secondary metabolites lacked a significant role in the ability of *P. protegens* DTU9.1 to colonize the soil-like environment axenically (Figure S2A). Importantly, wildtype *P. protegens* DTU9.1 produced detectable amounts of DAPG, pyoluteorin, and orfamide A after seven days of cultivation in the bead system, as verified using liquid chromatography coupled to high resolution-mass spectrometry (LC-HRMS) (Figure S2B). In this experimental setup, we did not detect other known secondary metabolites of *P. protegens*, such as pyrrolnitrin, pyoverdine and pyochelin.

We then investigated the effect of the three metabolites, DAPG, pyoluteorin, and orfamide A on the ability of *P. protegens* DTU9.1 to invade the SynCom. Based on the results of the initial MIC assays (Table 1), we hypothesized that the abundance of several SynCom members would be negatively affected by exposure to the three antimicrobial metabolites. After 24 hours of cultivation, all SynCom members were still detected (Figure 1A), whereas the amount of cultivable *P. protegens* DTU9.1 cells were 100-1000 fold lower than the SynCom members, likely owing to the low inoculum size of *P. protegens* DTU9.1 (see Methods). However, on the fourth day, *P. protegens* DTU9.1 had reached high cell numbers (≈10^8^ CFU/ml), which were maintained throughout the experiment and comparable to that observed in axenic cultivation. On the seventh day of incubation *P. protegens* DTU9.1 constituted the majority of the bacterial biomass regardless of variant (wildtype or mutants), while at least three of the four SynCom members remained detectable with our method of colony counting (Figure 1C). However, on the fourth day and onwards, colony forming units of *Pedobacter* sp. D749 were no longer detectable in the systems inoculated with the variants of *P. protegens* DTU9.1 (Figure 1B and 1C). Moreover, the control system without *Pseudomonas* allowed the persistence of *Pedobacter* sp. D749 at higher cell titers throughout the experiment, albeit with cell counts close to the detection limit at the seventh day (Figure 1C). Importantly, amplicon sequencing of the 16S rDNA gene confirmed the low presence of *Pedobacter sp.* D749 throughout the seven days in both the control system and the system with *P. protegens* DTU9.1 wildtype, despite cell numbers being below the countable detection limit (Figure S3). This shows that *Pedobacter* sp. D749 was not extinguished from the system upon invasion by *P. protegens* DTU9.1, but rather severely inhibited compared to the control system.

**Figure 1.**
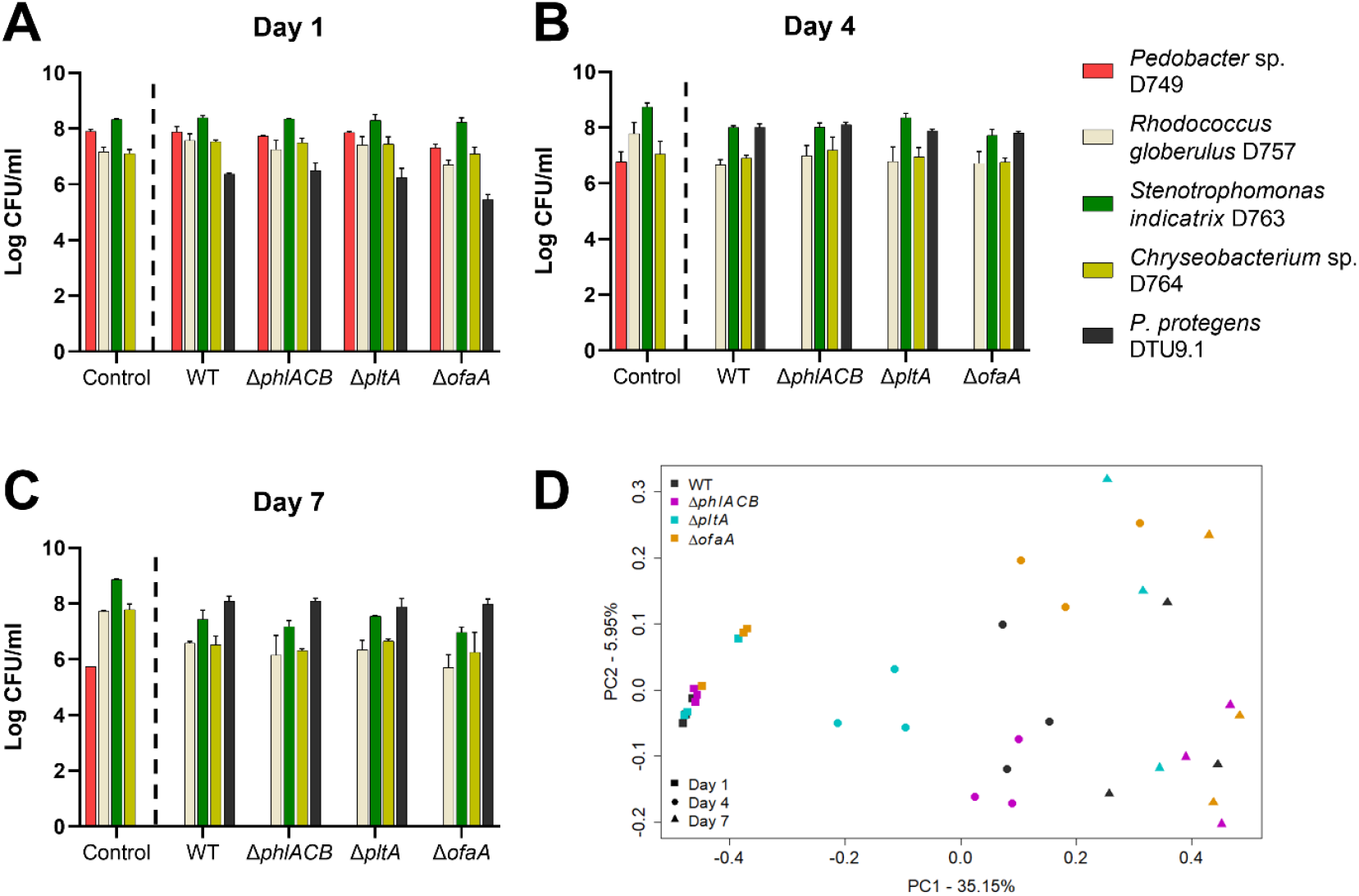
*P. protegens* DTU9.1 invades a four-membered synthetic microbial community in a soil-like environment. Abundance was determined as colony forming units in the hydrogel bead system of each SynCom member and the introduced *P. protegens* DTU9.1 variant; WT, Δ*phlACB*, Δ*pltA* and Δ*ofaA* on Day 1 (A), Day 4 (B) and Day 7 (C). The dotted line indicates the separation of data (right) used for the Principal Coordinate Analysis excluding the control system without *P. protegens*. Data was derived from three biological replicates. D) A Principal Coordinate Analysis (PCoA) was performed on CFU data of the systems inoculated with variants of *P. protegens* DTU9.1 to compare the effects of sampling time (symbols) and the variant of *P. protegens* DTU9.1 (color) on the bacterial composition in a soil-like environment.

The overall effect of DAPG, pyoluteorin and orfamide A on community composition over time was evaluated with a Principal Coordinate Analysis (PCoA) using Bray-Curtis distances (Figure 1D). The analysis was performed on non-logarithmic transformed CFU data of the bacterial abundance in the systems inoculated with variants of *P. protegens* DTU9.1 (WT, Δ*phlACB*, Δ*pltA*, and Δ*ofaA*) to allow visualization of more subtle changes. The analysis suggested that the major factor affecting abundance was the sampling time, as three separate clusters representing the three sampling times appeared across the first principal component, which explained most of the variance in the data (35.15%). An overall PERMANOVA using sampling time, genotypic variant of invading *P. protegens* DTU9.1, and their interaction as fixed effects further confirmed the significance of sampling time on the community composition (*P* = 9.99 · 10^-5^, R^2^ = 0.75). Measuring the growth rates of each SynCom member and the four variants of *P. protegens* DTU9.1 in liquid broth revealed that *Pseudomonas* grew significantly faster than each SynCom member (Figure S4), whereas *Pedobacter* sp. D749 had the slowest growth rate. This could explain the observed significant effect of sampling time on community composition, as visualized by the PCoA (Figure 1D).

### The secondary metabolome of *P. protegens* DTU9.1 is altered during cocultivation with a four-membered SynCom

Contrary to our initial hypothesis, the secondary metabolites (DAPG, pyoluteorin and orfamide A) seemed to play an insignificant role in the ability of *P. protegens* DTU9.1 to establish within or modify the SynCom composition. To investigate this observation further, we extracted metabolites from the hydrogel bead systems after seven days of incubation and analyzed the samples with LC-HRMS to verify production and persistence of the three secondary metabolites in question. During axenic cultivation, *P. protegens* DTU9.1 reached similar levels of colony forming units as compared to growth alongside the SynCom (Table 2). However, the concentrations of the secondary metabolites produced by *P. protegens* DTU9.1 differed markedly between the two systems. In the case of both DAPG and pyoluteorin, concentrations were elevated during cocultivation compared to axenic growth suggesting that *P. protegens* DTU9.1 responded to the presence of competing microorganisms by increasing the production of its antimicrobial secondary metabolites (*i.e.* DAPG and pyoluteorin). However, in the case of the cyclic lipopeptide, orfamide A, the concentration was significantly lower (*P* = 0.0002) when *P. protegens* DTU9.1 was cultivated with the SynCom compared to axenic growth (Table 2). This indicates that members of the synthetic community either inhibited production of orfamide A or affected the persistence of the metabolite over time as a potential mechanism of resistance.

**Table 2.** Concentration of secondary metabolites from *P. protegens* DTU9.1 after 7 days of growth in a soil-mimicking environment

|  | Log<sub>10</sub> CFU/ml<sup>a</sup><br>*Pseudomonas* | Secondary metabolite (µg/ml)<br>Mean ± Standard deviation |  |  |
| --- | --- | --- | --- | --- |
|  |  | DAPG | Pyoluteorin | Orfamide A |
| *P. protegens* DTU9.1 | 8.07 ± 0.11 | 3.96 ± 0.82 | 5.85 ± 0.56 | 25.58 ± 2.11 |
| *P. protegens* DTU9.1 + SynCom | 8.08 ± 0.15<sup>b</sup> | 9.16 ± 1.30 | 10.12 ± 2.19 | 3.87 ± 0.75 |
| Statistical significance<sup>c</sup> | n.s. | *P* = 0.0087 | *P* = 0.0556 | *P* = 0.0002 |

<sup>a</sup> CFU/ml of *P. protegens* DTU9.1 either grown alone or with the SynCom for seven days in the bead system.
<sup>b</sup> *P. protegens* DTU9.1 constituted approximately 75% of the total bacterial abundance in the microbial community.
<sup>c</sup> Significance of the difference in secondary metabolite concentration was analyzed by Student's *t*-test.

Notably, the average concentrations of DAPG and pyoluteorin observed after seven days of cultivation in the hydrogel bead system exceeded the minimum inhibitory concentration towards several SynCom members (Table 1). This could indicate that these species gained sufficient tolerance to sustain the levels of DAPG and pyoluteorin by growing in a 3D structure allowing for spatial distribution and biofilm formation, compared to planktonic growth in liquid broth. In the case of orfamide A, *P. protegens* DTU9.1 had the ability to produce 25.58 ± 2.11 µg/ml under axenic conditions (Table 2). However, during cocultivation with the SynCom the concentration of orfamide A was reduced to 3.87 ± 0.75 µg/ml, which is markedly below the MIC towards *R. globerulus* D757. This suggests that one or multiple SynCom members either inhibited production of orfamide A or secreted enzymes capable of inactivating orfamide A.

### Orfamide A is degraded during coculture of *P. protegens* DTU9.1 and SynCom in a soil-mimicking environment

To further investigate the fate of orfamide A (**1**) during coculture of *P. protegens* DTU9.1 and the SynCom, the MS data from the LC-HRMS analyses of metabolites was subjected to Global Natural Product Social (GNPS) molecular network analysis [22] to identify potential chemical relationships between features across the three systems (Figure S5). This revealed a distinct molecular family displaying the presence of orfamide A (*m/z* 1295.8509) during both axenic cultivation of *P. protegens* DTU9.1 and cocultivation with the SynCom (Figure 2A). Interestingly, the molecular family also contained features with related fragmentation patterns to orfamide A that only appeared during cocultivation of *P. protegens* DTU9.1 and the SynCom. One feature, *m/z* 1313.8542, corresponded to the mass of hydrolyzed orfamide A, with the addition of 18 Da (Figure 2A). Fragmentation analysis suggested hydrolysis of the ester bond (Figure 2B), which created a linearized congener of orfamide A (**2**) when *P. protegens* DTU9.1 was cultivated alongside the SynCom in the bead system (Figure 2D). Furthermore, potential degradation products of orfamide A were also present in the same molecular family (Figure 2A). The suspected degradation by-products were verified in the LC-HRMS data of the extracts. Differences in the retention times of each of the degradation by-products provided confirmation that these were not arising from in-source fragmentation (Figure S6). Fragmentation patterns were subsequently analyzed, confirming degradation products emanating from the hydrolyzed ester bond (Figure S7). A similar phenomenon was observed for orfamide B through the presence of degradation products (*m/z* 1113.7369, *m/z* 1000.6529, and *m/z* 887.5683 in Figure 2A). The remaining degradation products of orfamide B were present in concentrations too low to be selected for MS/MS, but were observed in the raw data (available from the raw data. See Methods). Taken together, these results demonstrate that one or multiple SynCom members are able to hydrolyze orfamide A (and orfamide B) from *P. protegens* DTU9.1 and degrade the linearized lipopeptide.

**Figure 2.**
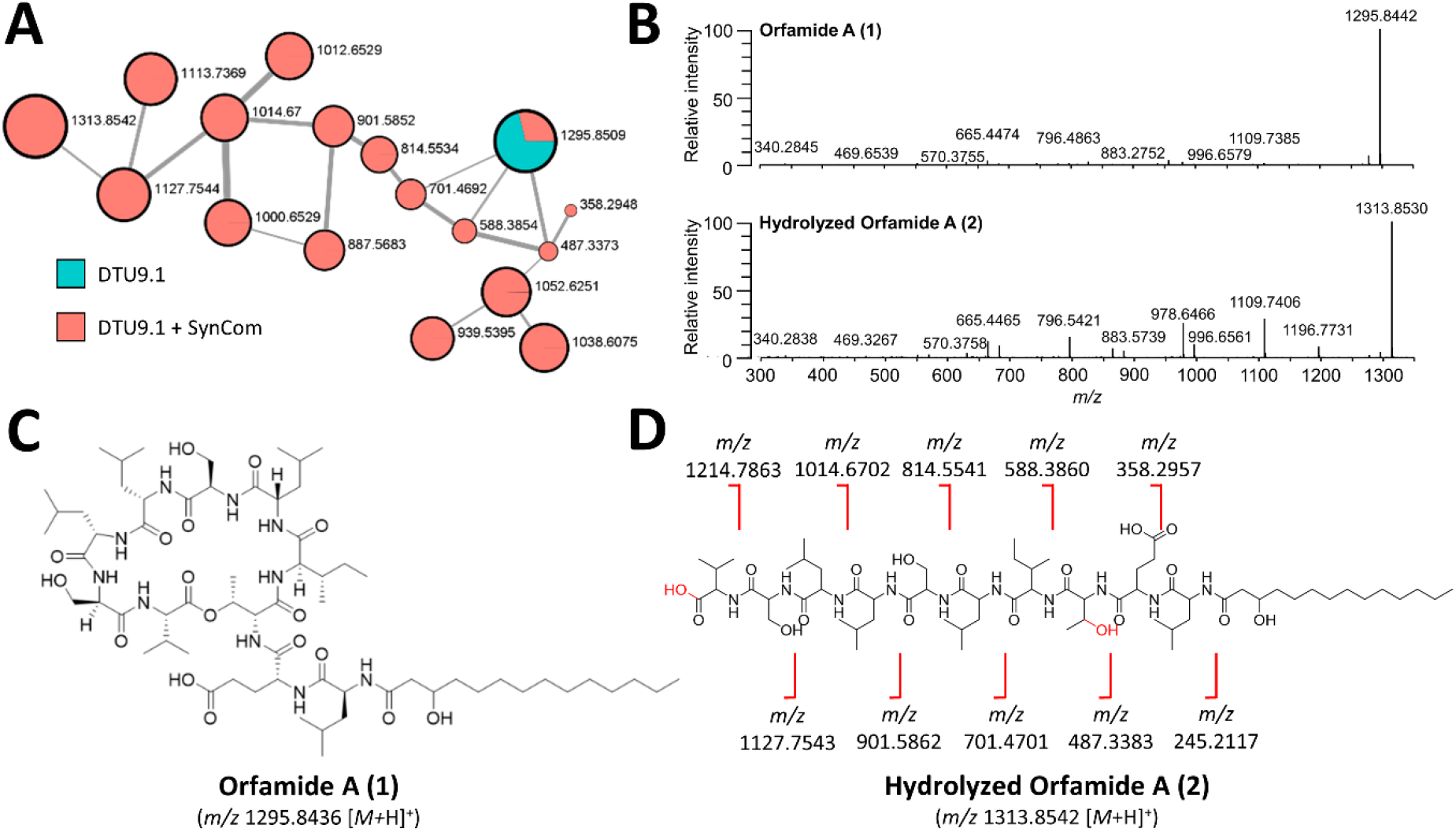
Orfamide A is degraded during coculture of *P. protegens* DTU9.1 and SynCom in a soil-mimicking environment. A) Molecular family of metabolites related to orfamide A derived from the molecular network (GNPS). Values next to nodes represent *m/*z-values and node size is scaled to metabolite mass. Colors inside nodes represent relative abundance in the system with axenically grown *P. protegens* DTU9.1 (Cyan) or the coculture between *P. protegens* DTU9.1 and the SynCom (Red). Edges between nodes represent molecular similarity based on cosine scores. B) Observed fragmentation patterns of 1 and 2 revealed by tandem mass spectrometry (MS/MS). C) Structure of orfamide A (1). D) Structure of hydrolyzed orfamide A (2) and the calculated masses of each degradation product.

### The fate of orfamide A is affected by a multi-species interaction involving *R. globerulus* D757 and *S. indicatrix* D763

To explore the degradation of orfamide A, we embarked upon dual-species interactions on agar surfaces to investigate if a single community member was responsible for linearization by hydrolysis and degradation. First, *P. protegens* DTU9.1 was cocultivated with each of the four SynCom members individually in mixed species colonies on 0.1x TSA. MS data of the metabolites extracted from an agar plug covering the entire bacterial colony revealed that the Gram-positive *R. globerulus* D757 was able to hydrolyze the ester bond of **1** yielding a linearized product, **2** (Figure 5A). Additionally, we monitored the interaction between *P. protegens* DTU9.1 and *R. globerulus* D757 by matrix-assisted laser desorption-ionization mass spectrometry imaging (MALDI-MSI) to validate that the linearization of orfamide A indeed occurs in the interface between the two bacteria. The two species were cultivated for 4 days on 0.1x TSA plate prior to matrix application and imaging. As displayed in Figure 3B, orfamide A (**1**) was secreted evenly around the *P. protegens* DTU9.1 colony, while the linearized product (**2**) was observed only in the interface between the two bacteria.

**Figure 3.**
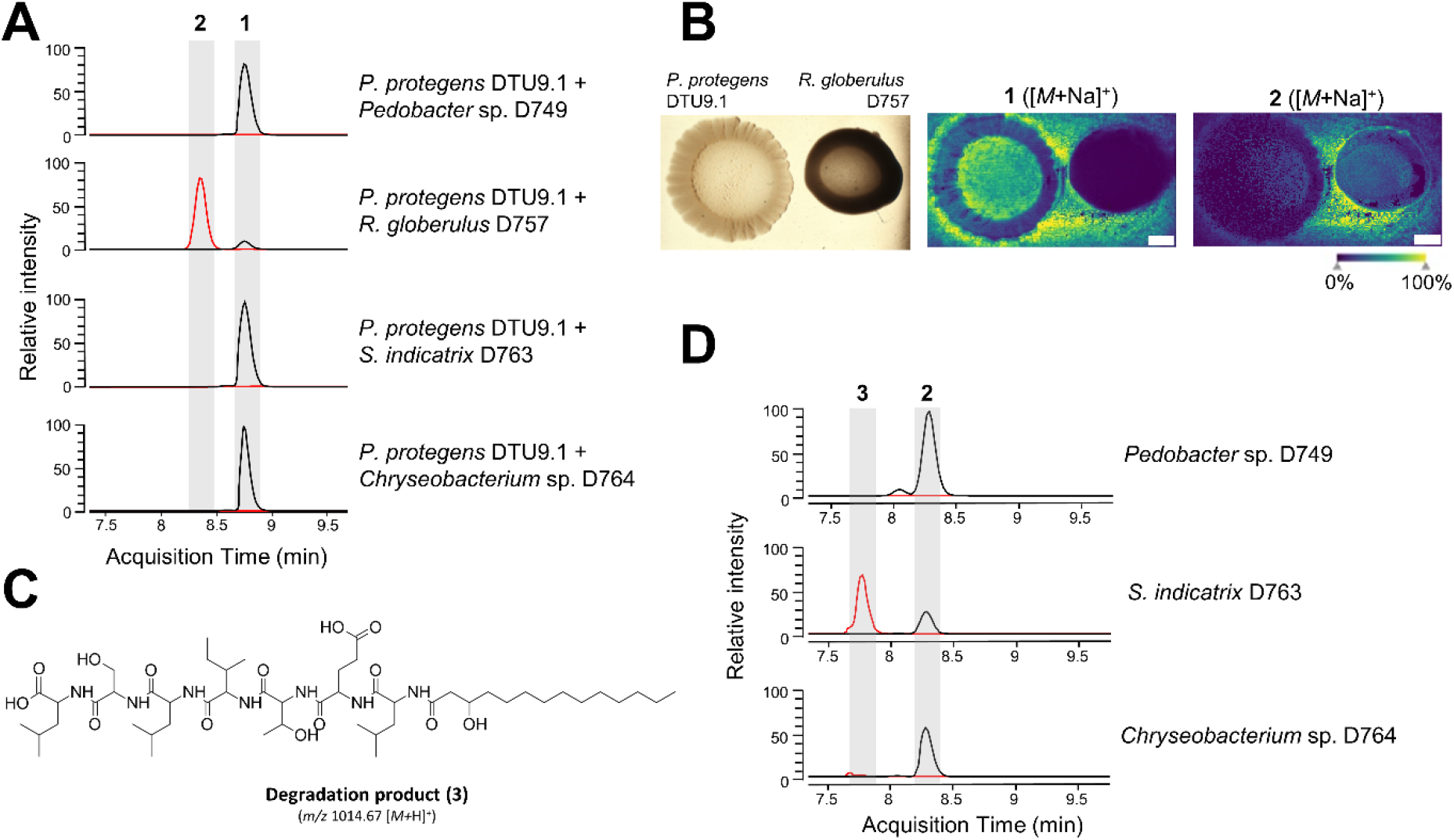
Orfamide A is hydrolyzed and degraded by separate members of the SynCom. A) Extracted ion chromatograms (EIC) for 1 (black, *m/z* 1295.8436 ± 5 ppm) and 2 (red, *m/z* 1313.8542 ± 5 ppm) detected in agar plugs from co-cultures of *P. protegens* DTU9.1 and the respective SynCom member on 0.1x TSA plates. B) MALDI imaging of colonies of *P. protegens* DTU9.1 and *R. globerulus* D757 showing hydrolysis of Orfamide A after 4 days of incubation. White scale bar represents 2 mm. C) Extracted ion chromatograms (EIC) for 2 (black, *m/z* 1313.8542 ± 5 ppm) and 3 (red, *m/z* 1014.6700 ± 5 ppm) detected in supernatants from cultures of *Pedobacter* sp. D749, *S. indicatrix* D763, and *Chryseobacterium* sp. D764 after 24 hours of cultivation.

In both assays we only observed hydrolysis of orfamide A, but no subsequent degradation. Thus, we hypothesized that one or more of the remaining three SynCom members (*Pedobacter* sp. D749, *S. indicatrix* D763, and *Chryseobacterium* sp. D764) were responsible for further degradation of the pre-hydrolyzed orfamide A (**2**). To investigate this hypothesis, we chemically hydrolyzed pure orfamide A by prolonged incubation in an alkaline solution (see Methods) and exposed the remaining three SynCom members individually to hydrolyzed orfamide A over 24 hours in liquid broth. Subsequent extraction of metabolites from the supernatants revealed that *S. indicatrix* D763 could degrade the hydrolyzed lipopeptide (Figure 3D). Here, we show the feature, *m/z* 1014.6700 (**3**), as an example of degradation. This feature corresponds to the mass of **2** excluding three amino acids from the C-terminal end (Figure 3C). However, we did also observe several of the larger degradation products as shown in Figure 2A (available from the raw data. See Methods). The lack of all degradation products as initially observed in the bead system could be explained by the reduced incubation time of 24 hours compared to seven days. Nonetheless, this demonstrates that *S. indicatrix* D763 can degrade **2** following prior hydrolysis of the ester bond in **1** by *R. globerulus* D757. Thus, the fate of orfamide A (**1**) in our experimental setup was affected by a sequential, multi-species interaction involving two co-isolated soil bacteria ultimately leading to complete degradation.

## Discussion

In this study, we utilized a four-membered bacterial SynCom cultivated in an artificial hydrogel soil environment to evaluate the contribution of secondary metabolites from a *P. protegens* strain on the ability of the *Pseudomonas* to invade and establish within the simplified microcosm. Although the genotypic complexity of our four-membered SynCom is far from a representation of the natural soil microbial diversity [23, 24], the simplified system applied in this study has allowed for the systematic analysis of community-level interactions affecting secondary metabolism in *P. protegens* DTU9.1. We demonstrated that *P. protegens* readily invaded and altered the community composition. Unexpectedly, mutants unable to produce these antibiotics invaded the community as efficiently as wildtype producers did and caused similar perturbations in the composition of the synthetic community (Figure 1). The absence of a clear antibiotic-mediated impact on the otherwise antibiotic-sensitive SynCom members (Table 2 and Figure S4) was not a result of low, non-toxic levels of antibiotics (Table 1). Similarly, previous studies reported the lack of advert effects of the biocontrol strain *P. protegens* CHA0 on indigenous microorganisms *in situ*, despite large fractions of subsequently isolated rhizobacteria displaying sensitivity towards antimicrobial metabolites produced by the biocontrol strain *in vitro* [11, 25]. This could suggest that sensitive bacteria either colonize distinct niches separated from the antibiotic-producing biocontrol strain *in situ* or that naturally co-occurring microorganisms sustain toxic levels of antimicrobial metabolites via community-level tolerance mechanisms, such as multi-species biofilm formation [26] or enzymatic inactivation of antimicrobial metabolites [27].

Microbial interactions among co-existing microorganisms have been studied extensively over the past decade to identify and understand the diverse means by which these microbes communicate and compete. Foster and Bell reported that interspecies competition among bacteria isolated from the same sample site was the far most dominant type of interaction [28]. This could suggest that bacteria have evolved intricate sensing mechanisms to respond to danger cues secreted by competing organisms [29, 30]. In our study, we found that the invading *P. protegens* DTU9.1 significantly increased the production of its antimicrobial metabolites, DAPG and pyoluteorin, during cocultivation with the four-membered SynCom (Table 1). This suggests that the biosynthesis of these metabolites is induced in *P. protegens* as a response to cues secreted by one or more competing members of the SynCom, which we have observed previously in the case of pyoluteorin biosynthesis [31]. However, as shown in Figure 1, the observed perturbations in community composition upon *Pseudomonas* invasion were not related to the production of antimicrobial secondary metabolites. Notably, the concentration of pyoluteorin exceeded the minimal inhibitory concentration towards *Pedobacter* sp. D749, *R. globerulus* D757, and *Chryseobacterium* sp. D764 on the seventh day of the experiment (Tables 1 and 2). However, this increase had no significant effect on abundance of either SynCom member compared to the system inoculated with the Δ*pltA* mutant. Although the mechanisms underlying the SynCom tolerance towards pyoluteorin and DAPG were not further pursued, we suggest that cultivation within the hydrogel bead system facilitate either formation of microbial biofilms, which alter the physiological state and antibiotic susceptibility of biofilm members [26, 32] or interspecies interaction that modulate bacterial growth properties and antibiotic tolerance of community members [33, 34].

Importantly, we observed a significant reduction in the concentration of the non-ribosomally synthesized cyclic lipopeptide, orfamide A, during cocultivation of *P. protegens* DTU9.1 and the four-membered SynCom. Lipopeptides from fluorescent *Pseudomonas* are versatile metabolites most notably known for their antimicrobial properties and involvement in bacterial motility [35, 36]. Particularly, Gram-positive bacteria have been associated with increased susceptibility towards lipopeptides, due to the lack of a protective cell wall [37], thus it is not surprising that some Gram-positive Actinobacteria have evolved resistance mechanisms towards lipopeptides involving enzymatic inactivation [38, 39]. In our study we discovered that the Gram-positive *R. globerulus* D757 inactivated orfamide A by hydrolysis of the thermodynamically sensitive ester bond connecting the macrocyclic ring. According to D’Costa and colleagues, hydrolysis of the ester bond is the most common resistance mechanism by which Actinobacteria enzymatically inactivated the cyclic lipopeptide, daptomycin [39]. While enzymatic inactivation of orfamide A is likely a resistance mechanism in *R. globerulus* D757, this biotransformation is also a prerequisite for subsequent degradation by *S. indicatrix* D763 (Figure 3D). Two recent studies have similarly demonstrated how biotransformation of *Pseudomonas*-produced cyclic lipopeptides can have dramatic effects on their chemical and ecological properties [20, 40]. Zhang et al. found that coculture between a *Pseudomonas* and a *Paenibacillus* strain led to the enzymatic modification of syringafactin, thus changing the chemical structure of the lipopeptide resulting in an amoebicidal byproduct [20]. Hermenau et al. demonstrated that Gram-positive *Mycetocola* strains could disarm the activity of the mushroom pathogen, *P. tolaasii*, by hydrolysis of the ester bonds in the two *Pseudomonas*-produced cyclic lipopeptides, tolaasin I and pseudodesmin A, thus preventing pathogenesis [40]. In our study, we identified catabolism of hydrolyzed orfamide A (Figure 2A and Figure 3D), which suggests that *S. indicatrix* D763 might utilize the hydrolyzed orfamide A as an alternative nutrient source. A prior study has demonstrated the ability of isolated soil bacteria to utilize penicillin as sole carbon source to support growth by enzymatically catabolizing the antibiotic [41]. Further investigation of the impact of orfamide A catabolism will be the subject for future studies.

It has previously been shown that *Pseudomonas*-produced cyclic lipopeptides are rapidly degraded when added exogenously to non-sterile soil yet remain stable in sterilized soil, which clearly suggests that unknown members of the indigenous soil microbiome possess the ability to transform and/or degrade metabolites with activities relevant for biocontrol [42]. Our discovery and characterization of community-level inactivation and degradation of orfamide A involving two co-occurring soil bacteria in our SynCom provides a mechanistic explanation for this observation. Although the prevalence of such processes in different soil communities is currently unknown, we suggest that biotransformation processes of cyclic lipopeptides and other biocontrol metabolites [43] may contribute to variations in efficiency in biocontrol applications. Collectively, our results illustrate the usefulness of synthetic communities to systematically investigate how microbial communities respond to antibiotics to enhance their resilience towards microbial invasion, and to identify processes that determines the turnover and ‘fate’ of biocontrol metabolites. Improved knowledge of potential constraints in efficient biocontrol is a prerequisite for development of efficient biocontrol products or for development of measures to counteract the community processes responsible for modification and degradation of biocontrol metabolites.

## Methods

### Microorganisms and cultivation

Plasmid cloning was performed in *Escherichia coli* CC118-*λpir*. Cells were cultured in lysogeny broth (LB; Lennox, Merck, St. Louis, MO, USA) with appropriate antibiotics. The antibiotic concentration used was 10 µg/ml for chloramphenicol, and 8 µg/ml and 50 µg/ml for tetracycline for *E. coli* and *P. protegens*, respectively. *E. coli* CC118 λ*pir* was cultured by inoculating a single colony in 5 ml LB broth and incubating overnight at 37° C with shaking (200 rpm). *P. protegens* DTU9.1 and members of the semi-synthetic community were cultured by inoculating a single colony in 5 ml LB broth and incubating overnight at 30° C with shaking (200 rpm). The community members include *Pedobacter* sp. D749 (Accession: CP079218), *R. globerulus* D757 (Accession: CP079698), *S. indicatrix* D763 (Accession: CP079106), and *Chryseobacterium* sp. D764 (Accession: CP079219) [19].

### Generation of secondary metabolite deficient mutants

To generate mutants in *P. protegens* DTU9.1 incapable of synthesizing DAPG, pyoluteorin and orfamide A, genes required for biosynthesis were deleted by allelic replacement according to Hmelo et al. [44]. Primers used for cloning and verification are summarized in Supplementary Table 1. In short, DNA fragments directly upstream and directly downstream of *the gene of interest* were PCR amplified and subsequently joined by splicing-by-overlap extension PCR with XbaI and SacI overhangs. The purified PCR product was restriction-digested and inserted in pNJ1 [45]. The resulting plasmid was mobilized into *P. protegens* DTU9.1 via triparental mating with *E. coli* HB101 harboring the helper plasmid pRK600. Merodiploid transconjugants were initially selected on *Pseudomonas* Isolation Agar (PIA, Merck) supplemented with 50 µg/ml tetracycline. A second selection was performed on NSLB agar (10 g/l tryptone, 5 g/l yeast extract, 15 g/l Bacto agar) with 15% v/v sucrose. Candidates for successful deletion were confirmed by PCR and verified by Sanger sequencing at Eurofins Genomics.

### Integration of *P. protegens* DTU9.1 in a semi-synthetic microbial community

The effect of introducing *P. protegens* DTU9.1 and the above-mentioned secondary metabolite-deficient mutants into a semi-synthetic bacterial community was investigated in an artificial soil medium composed of spherical hydrogel beads. The beads were prepared according to Ma et al. [18]. In short, a polymer solution was prepared as a 4:1 mixture of 9.6 g/l gellan gum (Phytagel^TM^, Sigma) and 2.4 g/l sodium alginate (Sigma) dissolved in distilled water. Spherical beads with a diameter of approximately 3-4 mm were formed by dropping polymer solution into a cross-linker solution containing 20 g l^-1^ CaCl_2_ with a 10 ml syringe. Then, the beads were soaked in 0.1x TSB (Sigma) for 1 hour followed by sieving the beads to remove residual TSB medium. Finally, 20 ml beads were transferred to 50 ml Falcon tubes. Cultures of the four community members and *P. protegens* WT Δ*phlACB*, Δ*pltA* and Δ*ofaA* were grown overnight (O/N). The optical density at 600 nm (OD_600_) of *Pedobacter* and *Rhodococcus* was set to 2.0, for *Stenotrophomonas* and *Chryseobacterium* it was set to 0.1 and for *P. protegens* DTU9.1 and mutants it was set to 0.001. Bacterial inoculation suspensions were prepared by mixing equal volumes in a total volume of 2 ml. Lastly, the prepared beads were inoculated with the 2 ml bacterial suspension. Inoculated bead systems were incubated static at RT and samples collected after 1, 4 and 7 days. Sampling was performed by briefly shaking the bead systems followed by extracting approximately 1 ml beads into new 15 ml Falcon tubes. Extracted beads were subsequently diluted in 0.9% (w/v) NaCl according to their weight to normalize the amount of bacterial cells. The tubes were shaken on a vortex for 10 minutes at maximum speed to disrupt the hydrogel beads. After vortexing, dilutions were spread on 0.1x TSA plates and incubated at RT for 48 hours before counting CFU/ml. The remaining liquid (approx. 5 ml) of the processed samples were saved for chemical detection of secondary metabolites.

### Minimal Inhibitory Concentration (MIC) assay

To determine the susceptibility of each SynCom member towards the three *Pseudomonas*-produced metabolites (DAPG, pyoluteorin and orfamide A) MIC assays were conducted. SynCom members were cultured in four biological replicates in LB O/N. Cells were washed twice in 0.9% NaCl. A clear 96-well flat-bottom microplate (Greiner Bio-One) was prepared with 200 µl 0.1x TSB per well inoculated with bacteria to an initial OD_600_ of 0.01 and appropriate serial dilutions of metabolites. The microplate was covered with semi-permeable membrane (Breathe-Easy, Merck) and incubated at room temperature with 600 rpm shaking for 24 hours, followed by MIC-value determination.

### Detection of secondary metabolites with LC-HRMS

To extract secondary metabolites in the hydrogel bead samples and supernatants of O/N cultures, an equal volume of ethyl acetate was added to the samples followed by shaking the tubes briefly. For extraction of metabolites from 0.1x TSA plates, an agar plug covering entire bacterial colonies (approx. 6mm diameter) was suspended in 1 ml isopropanol:ethyl acetate (1:3 v/v) with 1% formic acid and shaken briefly. For both types of extractions, tubes were subsequently centrifuged for 3 minutes at 4650 x g and the top ethyl acetate layer was transferred to new tubes. Extracts were then evaporated under N_2_ O/N. The dried extracts were re-suspended in 200 µl methanol (MeOH) and centrifuged for 3 minutes at 4650 x g. The supernatant was transferred to HPLC vials and subjected to ultra high-performance liquid chromatography electrospray ionization time-of-flight mass spectrometry (UHPLC-HRMS) analysis.

LC-HRMS was performed on an Agilent Infinity 1290 UHPLC system. Liquid chromatography of 1 µl or 5 µl extract was performed using an Agilent Poroshell 120 phenyl-C_6_column (2.1 × 150 mm, 1.9 μm) at 60 °C using CH_3_CN and H_2_O, both containing 20 mM formic acid. Initially, a linear gradient of 10% CH_3_CN/H_2_O to 100% CH_3_CN over 10 min was employed, followed by isocratic elution of 100% CH_3_CN for 2 min. Then, the gradient was returned to 10% CH_3_CN/H_2_O in 0.1 min and finally isocratic condition of 10% CH_3_CN/H_2_O for 1.9 min, all at a flow rate of 0.35 min/ml. HRMS data was recorded in positive ionization on an Agilent 6545 QTOF MS equipped with an Agilent Dual Jet Stream electrospray ion (ESI) source with a drying gas temperature of 250°C, drying gas flow of 8 min/l, sheath gas temperature of 300°C and sheath gas flow of 12 min/l. Capillary voltage was 4000 V and nozzle voltage was set to 500 V. Fragmentation data was collected using auto MS/MS at three collision energies (10, 20, 40 eV). The HRMS data was processed and analyzed using Agilent MassHunter Qualitative Analysis B.07.00. HPLC grade solvents (VWR Chemicals) were used for extractions while LCMS grade solvents (VWR Chemicals) were used for LCMS.

### GNPS molecular networking

A molecular network was created using the Feature-Based Molecular Networking [46] workflow on GNPS (https://gnps.ucsd.edu, [22]). The workflow run can be found at this link: https://gnps.ucsd.edu/ProteoSAFe/status.jsp?task=1ad207802221433ca5431d22f2638d0e. Raw data was processed using MZmine2.53 [47]. Data was filtered by removing all MS/MS fragment ions within +/− 17 Da of the precursor m/z. MS/MS spectra were window filtered by choosing only the top 6 fragment ions in the +/− 50 Da window throughout the spectrum. Additional settings include: precursor ion mass tolerance was set to 0.05 Da, MS/MS fragment ion tolerance to 0.05 Da, and edges were filtered to have a cosine score above 0.7 and more than 10 matched peaks. The spectra in the network were then searched against GNPS spectral libraries [22, 48]. The library spectra were filtered in the same manner as the input data. All matches kept between network spectra and library spectra were required to have a score above 0.7 and at least 6 matched peaks. The DEREPLICATOR was used to annotate MS/MS spectra [49]. The molecular networks were visualized using Cytoscape 3.8.2 [50].

### MALDI Sample preparation

Bacterial strains were cultured on 10 ml 0.1x TSA plates at 22°C. After 4 days of incubation, the microbial colony and surrounding agar was sectioned and mounted on an IntelliSlides conductive tin oxide glass slide (Bruker). The sample was covered with matrix by spraying 1.75 ml of a matrix solution in a nitrogen atmosphere. The matrix solution was 2,5-dihydrobenzoic acid (DHB) of 20 mg/ml concentration in ACN/MeOH/H_2_O (70:25:5, v/v/v) according to [51].

### MALDI Mass Spectrometry Imaging (MSI)

Samples were dried in a desiccator overnight prior to MSI measurement. The samples were then subjected to timsTOF flex mass spectrometer (Bruker) for MALDI-IMS acquisition. Calibration was done using red phosphorus. The samples were run in positive MS scan mode with 100 µm raster width and a *m/z* range of 100-2000. Briefly, a photograph of the colonies was loaded onto Fleximaging software, three teach points were selected to align the background image with the sample slide, measurement regions were defined, and the automatic run mode was then employed. The settings in the timsControl were as follow: Laser: imaging 100 µm, Power Boost 3.0%, scan range 26 µm in the XY interval, and laser power 90%; Tune: Funnel 1 RF 300 Vpp, Funnel 2 RF 300 Vpp, Multipole RF 300 Vpp, isCID 0 eV, Deflection Delta 70 V, MALDI plate offset 100 V, quadrupole ion energy 5 eV, quadrupole loss mass 100 *m/z*, collision energy 10 eV, focus pre TOF transfer time 75 µs, pre-pulse storage 8 µs. After data acquisition, the data was analyzed using SCiLS software.

### Chemical hydrolysis of orfamide A

Pure orfamide A (Cayman, United States) was chemically linearized by hydrolysis by mixing 100 µl (0.386 µmol, suspended in methanol) and 193 µL 0.1 M aqueous LiOH (1.93 µmol, 5 equimolar). The solution was stirred at room temperature for 21 h. The reaction mixture was quenched by addition of 29.3 µL 1 M HCl. This led to the formation of a white precipitate, which was re-dissolved by addition of 677.7 µl methanol. Complete hydrolysis was verified by LC-HRMS.

### Statistics

Multivariate analysis of community composition was performed using PERMANOVA on Bray-Curtis distances and the model formulation Y ∼ Time + Variant + Time:Variant. Follow-up PERMANOVAs were performed on each time point with only Variant as the dependent variable. A univariate comparison of CFU counts and metabolite concentrations in SynCom versus axenic culture of *P. protegens* DTU9.1 was carried out using Student’s *t*-tests assuming equal variance.

### Data availability

LC-HRMS data has been deposited at MassIVE with the identifier, MSV000092145. MALDI-MSI has been uploaded to Metaspace (https://metaspace2020.eu/project/Hansen-2023). Demultiplexed 16S rDNA sequencing reads were uploaded to NCBI SRA database under BioProject number PRJNA983551.

## Acknowledgements

We thank the members of the Center for Microbial Secondary Metabolites (CeMiSt) for general scientific discussions and advice.

