## Supplementary for "Resistance towards and biotransformation of *Pseudomonas*-produced secondary metabolites during community invasion"

### Supplementary methods

#### Measurement of growth rates

To calculate growth rates of each SynCom member, as well as the four variants of *P. protegens* DTU9.1 (WT, Δ*phlACB*, Δ*pltA*, Δ*ofaA*) all bacteria were cultured in four biological replicates in lysogeny broth (LB) overnight (O/N). Cells were washed twice in 0.9% NaCl. A clear 96-well flat-bottom microplate (Greiner Bio-One) was prepared with 200 µl 0.1x Tryptic Soy Broth (TSB, Sigma) per well inoculated with bacteria to an initial optical density at 600 nm ($OD_{600}$) of 0.01. The microplate was covered with semi-permeable membrane (Breathe-Easy, Merck) and incubated in a Cytation5 microplate reader (BioTek Instruments) at 22°C and continuous shake. Measurements of optical density at $OD_{600}$ were taken every 10 minutes for 24 hours. Growth rates were estimated with a linear regression during exponential growth and compared statistically with ANOVA followed by a Tukey *post-hoc* analysis.

#### DNA extraction from hydrogel bead systems and 16S rDNA amplicon sequencing

DNA from the hydrogel bead systems were extracted after 1, 4 and 7 days with a DNAeasy PowerSoil kit (Qiagen) following the manufacturer's instructions. The V3-V4 region of the 16S rDNA gene was amplified in a 25 µl PCR reaction (5.9 µl Sigma Water, 12.5 µl 2x TEMPase, 0.8 µl 10 µM barcoded 16S-341F, 0.8 µl 10 µM barcoded 16S-805R [1], and 5 µl template DNA). The program for the PCR reaction was as follows: (i) 15 min at 95°C, (ii) 30 cycles of 30 sec at 95°C, 30 sec at 60°C, and 30 sec at 72°C, and (iii) 5 min at 72°C. Amplicons were purified with the Agencourt AMPure XP kit (Beckman Coulter) and eluted in 10 mM Tris-HCl (pH = 8.5). Following purification, the amplicons were pooled in equimolar concentrations and shipped to Novogene (Cambridge, United Kingdom) for250 PE sequencing on an Illumina NovaSeq 6000 platform with 3 Gb raw data per sample. The raw reads were demultiplexed using cutadapt v3.7 [2]. We used *DADA2* 1.16 [3] in *R* 4.0.2 to denoise, join reads and perform downstream analyses with the scripts generated by Henriksen et al. [4].

##### Inhibition assays

The *in vitro* inhibitory effect of *P. protegens* DTU9.1 and the secondary metabolite deficient mutants was examined with agar plate inhibition assays. The members of the synthetic microbial community were the target bacteria. For the antibacterial inhibition assay, cultures of *P. protegens* DTU9.1 WT, Δ*phlACB*, Δ*pltA* and Δ*ofaA* were grown O/N. The cultures were washed twice in 0.9% (w/v) NaCl at 10,000 rpm for 1 minute followed by measuring the optical density at $OD_{600}$. The optical density was set to 1.0 and 10µl bacterial suspension was spotted on 0.1x Tryptic Soy Agar (TSA, Sigma). The plates were incubated at RT for 24 hours to allow for secondary metabolite production. O/N cultures of the target bacteria were prepared in LB medium. After 24 hours of incubation, the *Pseudomonas* on the 0.1x TSA plates were killed with chloroform vapors for 30 minutes, followed by evaporation of residual chloroform in a fume hood for another 30 minutes. The $OD_{600}$ of the target bacteria was set to 2.0 and 500µl was added to 6ml 0.1x TSB containing 0.3% agar pre-heated to 42°C. The soft agar was spread evenly on top of the 0.1x TSA plates with killed *Pseudomonas* and incubated at RT for 48 hours followed by examining the inhibition zones.

Supplementary Table 1

| **Name** | **Sequence**[a,b,c] | **Note** |
|---|---|---|
| ***phlACB* deletion** | | |
| Up_F$_{phlA}$ | 5'- atcccgtctagaCAGAGATTTCGCAGTAAAAAG | Amplification of homology region upstream of *phlA* |
| Up-R$_{phlA}$ | 5'- **catcgacgatttccgaagcgatcac**TTTCCTCTTGA TTCCATTCTTTTC | Amplification of homology region upstream of *phlA* |
| Down_F$_{phlB}$ | 5'- GTGATCGCTTCGGAAATCG | Amplification of homology region downstream of *phlB* |
| Down_R$_{phlB}$ | 5'- atccgggagctcACAACGAGGAGAATTCCAG | Amplification of homology region downstream of *phlB* |
| *phlACB*-del_fw | 5'- GTGCGAGTTCAATCATCTGG | Verification of *phlACB* deletion |
| *phlACB*-del_rev | 5'- CTCTCGTAGTTGAGCCGTTC | Verification of *phlACB* deletion |
| ***pltA* deletion** | | |
| Up_F$_{pltA}$ | 5'-atcccgtctagaAGCGCCTTCATTCCTAAATC | Amplification of homology region upstream of *pltA* |
| Up-R$_{pltA}$ | 5'- **gatgccaaagtaatgcgcatcgaag**TGCCCACT CCCTGTTAGGC | Amplification of homology region upstream of *pltA* |
| Down_F$_{pltA}$ | 5'- CTTCGATGCGCATTACTTTG | Amplification of homology region downstream of *pltA* |
| Down_R$_{pltA}$ | 5'- atccgggagctcCTGTCCAGCGAAGAGAGTT | Amplification of homology region downstream of *pltA* |
| *pltA*-del_fw | 5'- GTTCTTTGCATGTTCGAGAAAGAGCAG | Verification of *pltA* deletion |
| *pltA*-del_rev | 5'- GGGAACGCTTCCAGTCCACC | Verification of *pltA* deletion |
| ***ofaA* deletion** | | |
| Up_F$_{ofaA}$ | 5'- atcccgtctagaCTGCGACAGGCTCTTGAG AAAC | Amplification of homology region upstream of *ofaA* |
| Up-R$_{ofaA}$ | 5'- **ccggcgcggggaaaagcacttgcagcggcc**TTCA TGCGCGCCCCC | Amplification of homology region upstream of *ofaA* |
| Down_F$_{ofaA}$ | 5'- GGCCGCTGCAAGTGCTTTTC | Amplification of homology region downstream of *ofaA* |
| Down_R$_{ofaA}$ | 5'- atccgggagctcCCAGATGGTTGCGAA ACGGTAC | Amplification of homology region downstream of *ofaA* |
| *ofaA*-del_fw | 5'- CGCATTGATCAGCCTCGTGAC | Verification of *ofaA* deletion |
| *ofaA*-del_rev | 5'- GTAGTTGAGCATGGCGCTGAAC | Verification of *ofaA* deletion |
| **16S rDNA amplification** | | |
| 16S-341F | 5'- CCTACGGGNGGCWGCAG | Amplification of the V3-V4 region of the 16S rDNA gene |
| 16S-805R | 5'- GACTACHVGGGTATCTAATCC | Amplification of the V3-V4 region of the 16S rDNA gene |

*a.* CAPITAL letters represent the priming part

*b.* Underlined characters represent restriction sites attached as primer overhang

*c.* **Bold** characters represent the first 25 nucleotides downstream of the region targeted for deletion to achieve successful overlap-extension PCR

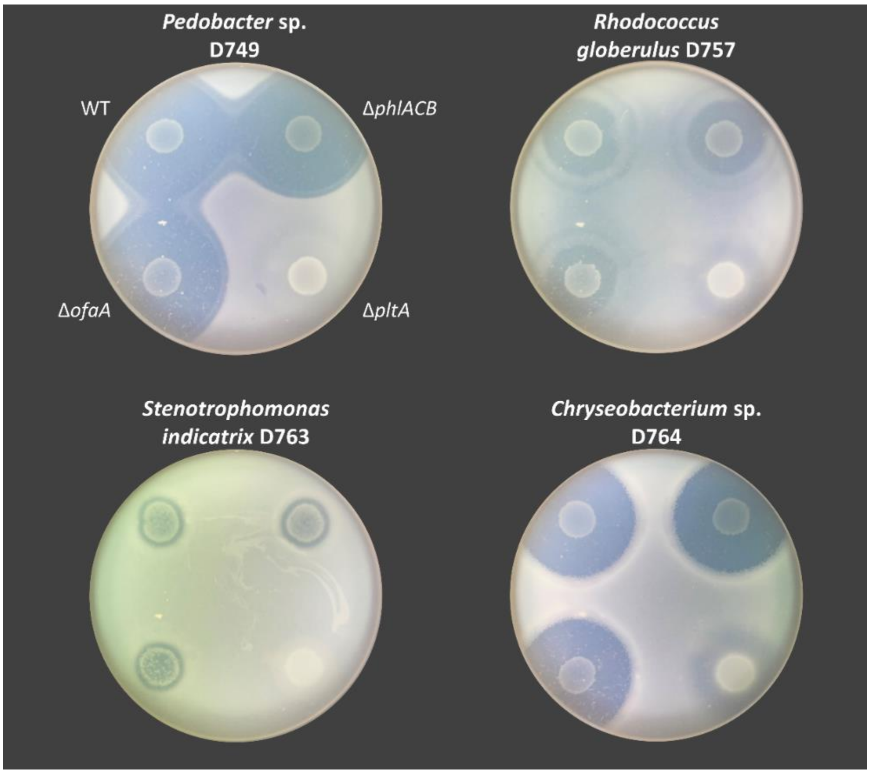


Figure S1 | **Pyoluteorin from *P. protegens* DTU9.1 is bactericidal towards all members of a synthetic microbial community.** Inhibition assay analyzing the *in vitro* bactericidal effect of DAPG, pyoluteorin and orfamide A on the four SynCom members: *Pedobacter* sp. D749, *R. globerulus* sp. D757, *S. indicatrix* D763, and *Chryseobacterium* sp. D764

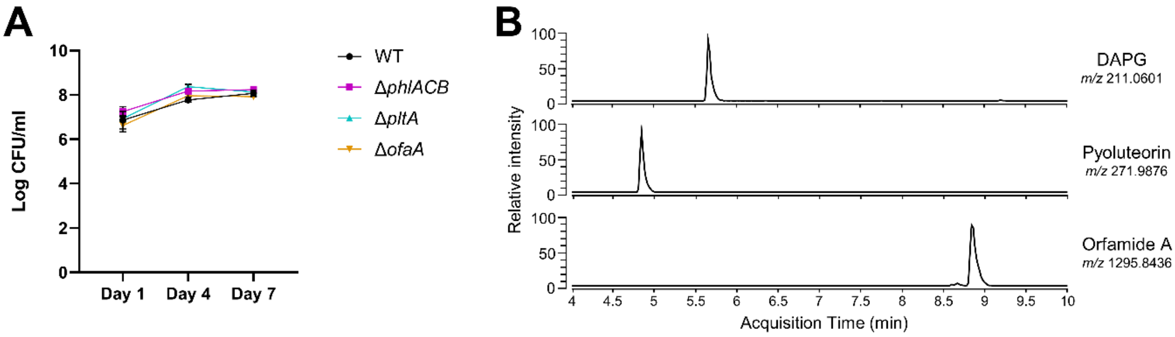


Figure S2 | ***P. protegens* DTU9.1 thrives in a soil-mimicking environment. A)** Growth curves of *P. protegens* DTU9.1 WT, Δ*phlACB*, Δ*pltA* and Δ*ofaA* in the hydrogel bead system. Samples were collected after 1, 4 and 7 days from three biological replicates. **B)** Extracted ion chromatograms (EIC) for DAPG (*m/z* 211.0601 ± 5 ppm), Pyoluteorin (*m/z* 271.9876 ± 5 ppm) and orfamide A (*m/z* 1295.8436 ± 5 ppm) confirm the production of these secondary metabolites after 7 days of growth of *P. protegens* DTU9.1 WT in the hydrogel bead system.

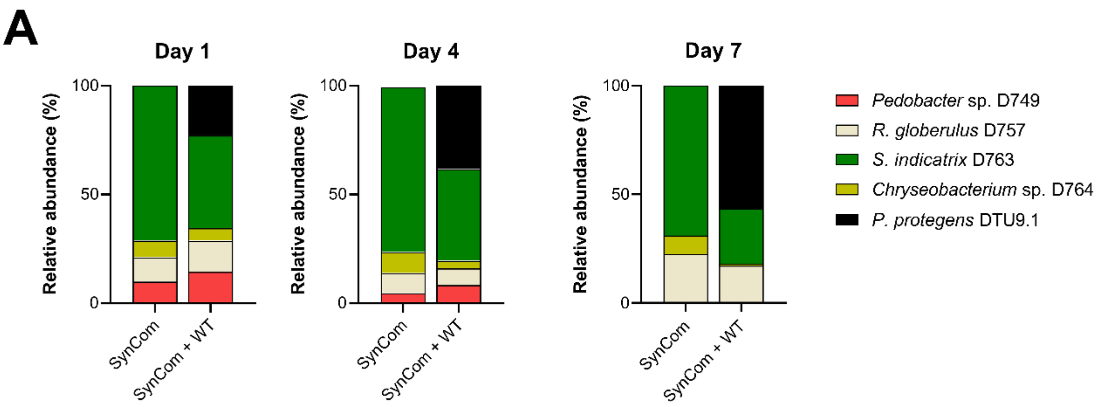


**B**

| | **Relative abundances in percentages** (Mean ± SD) | | | | | |
|---|---|---|---|---|---|---|
| | **Day 1** | | **Day 4** | | **Day 7** | |
| | SynCom | SynCom + WT | SynCom | SynCom + WT | SynCom | SynCom + WT |
| *Pedobacter* sp. D749 | 9.95 ± 3.18 | 14.44 ± 0.74 | 4.77 ± 1.50 | 8.40 ± 0.83 | 0.04 ± 0.02 | 0.38 ± 0.19 |
| *R. globerulus* D757 | 11.12 ± 1.77 | 14.29 ± 0.48 | 9.50 ± 1.29 | 7.60 ± 0.39 | 22.53 ± 4.16 | 16.79 ± 3.49 |
| *S. indicatrix* D763 | 71.19 ± 5.36 | 42.62 ± 2.32 | 76.02 ± 8.67 | 42.17 ± 2.44 | 68.96 ± 8.94 | 25.26 ± 2.77 |
| *Chryseobacterium* sp. D764 | 7.74 ± 1.64 | 5.80 ± 1.28 | 9.71 ± 6.06 | 3.57 ± 0.91 | 8.47 ± 4.77 | 0.94 ± 0.19 |
| *P. protegens* DTU9.1 | | 22.84 ± 2.49 | | 38.26 ± 2.66 | | 56.64 ± 3.91 |

Figure S3 | **Amplicon sequencing of the 16S rDNA gene of the SynCom with and without *P. protegens* DTU9.1 invasion. A)** Relative abundances of each bacterial species after 1, 4 and 7 days of cultivation within a hydrogel bead system. Number of reads were normalized to the presence of 16S rDNA genes per genome. Stacked bars represent the mean of three biological replicates. **B)** Relative abundances in percentages plotted in **A)**. Values represent mean and standard deviation from the three biological replicates.

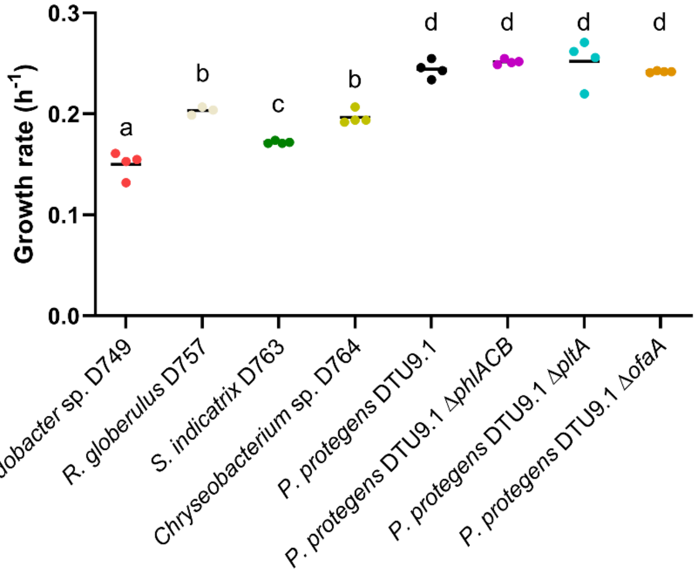


Figure S4 | ***P. protegens* DTU9.1 grows significantly faster than the four SynCom members.** Growth rates of *Pedobacter* sp. D749, *R. globerulus* sp. D757, *S. indicatrix* D763, and *Chryseobacterium* sp. D764., as well as *P. protegens* DTU9.1 WT, Δ*phlACB*, Δ*pltA*, and Δ*ofaA* in 0.1x TSB liquid broth. Data was derived from 4 biological replicates. Letters above indicate treatments significantly different from one another, as determined by a Tukey-test ($P < 0.05$).

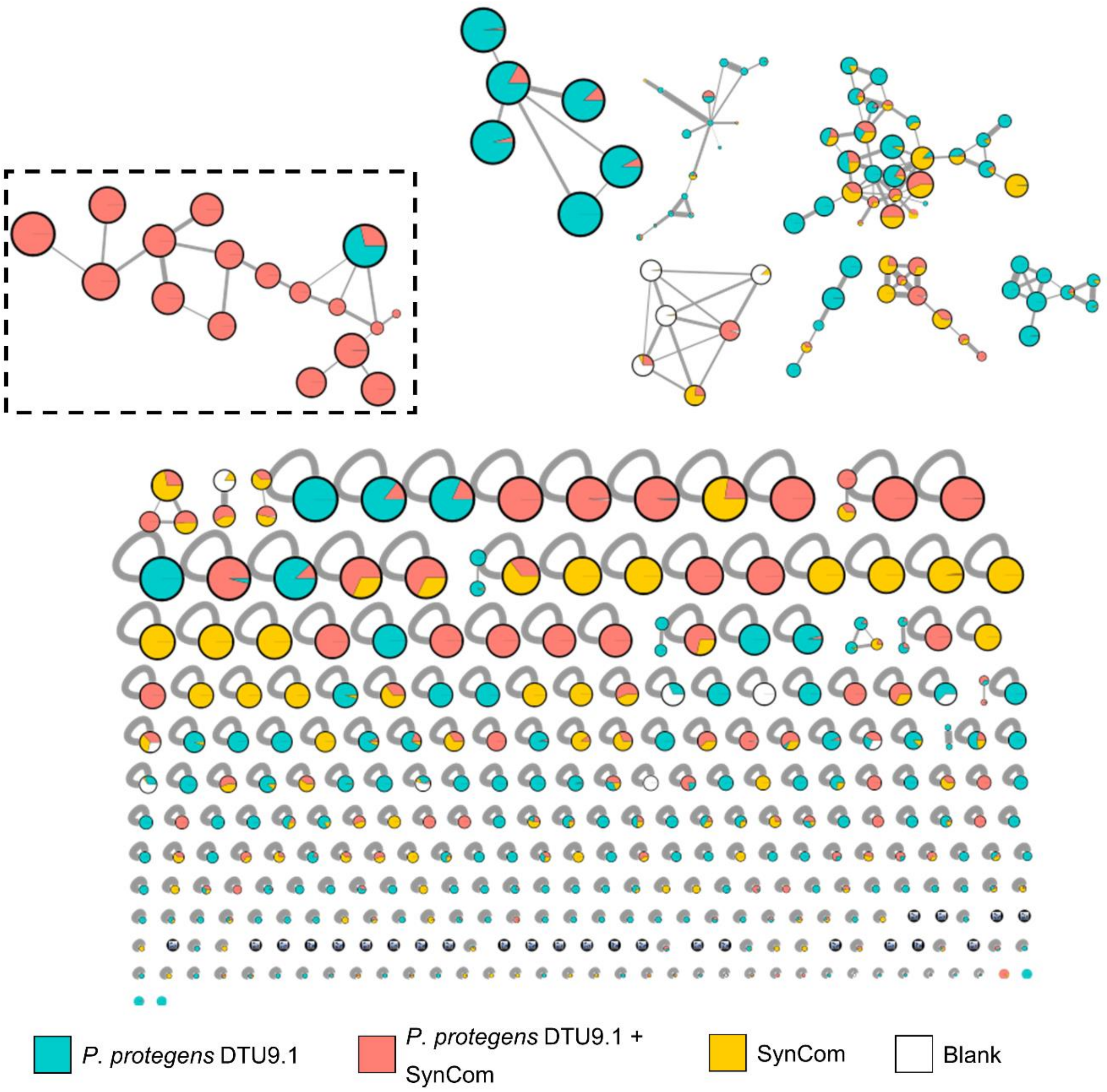


Figure S5 | **Global Natural Products Social network analysis of secondary metabolites extracted from bead systems.** Complete GNPS network on the MS data of the extracts from bead systems grown with monoculture of *P. protegens* DTU9.1 (Cyan), coculture between *P. protegens* DTU9.1 and SynCom (Red), or SynCom by itself (Yellow). White-colored nodes represent instrument blank noise. Nodes represent individual metabolites and the internal pie diagrams display relative abundance. The size of the nodes is scaled to metabolite mass. Edges between nodes represent the similarities between metabolites based on cosine score. The highlighted molecular family contain orfamide A and its respective degradation products identified during coculture.

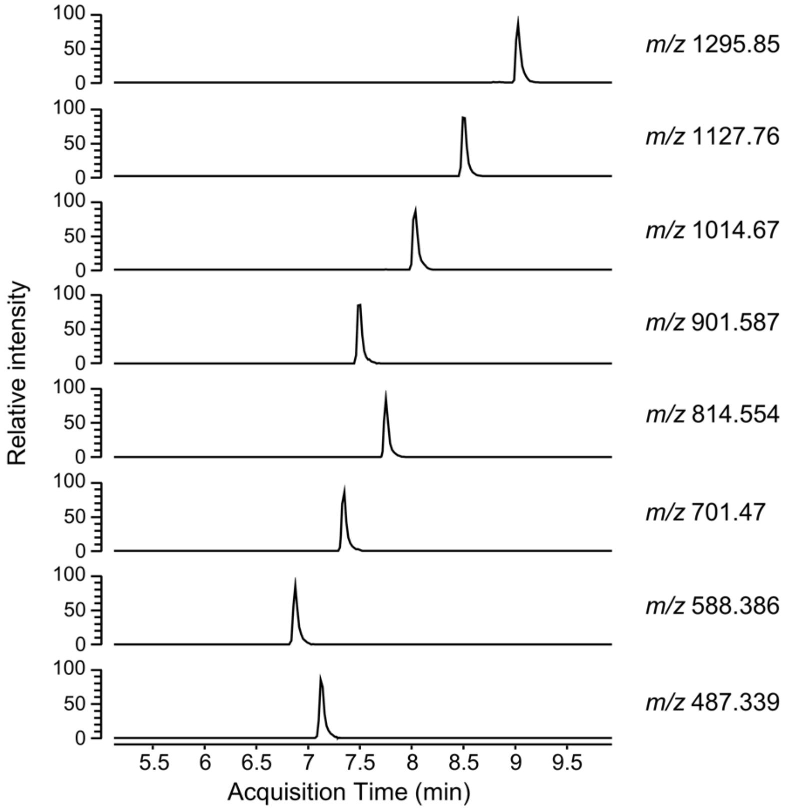


Figure S6 | **Extracted ion chromatograms confirmed the presence of degradation products.** Extracted ion chromatograms (EIC) of orfamide A and the observed degradation products after 7 days of cocultivation between *P. protegens* DTU9.1 and the SynCom in the hydrogel bead system. Each degradation product has a different retention time, thus denying the possibility of them being products of in-source fragmentation.

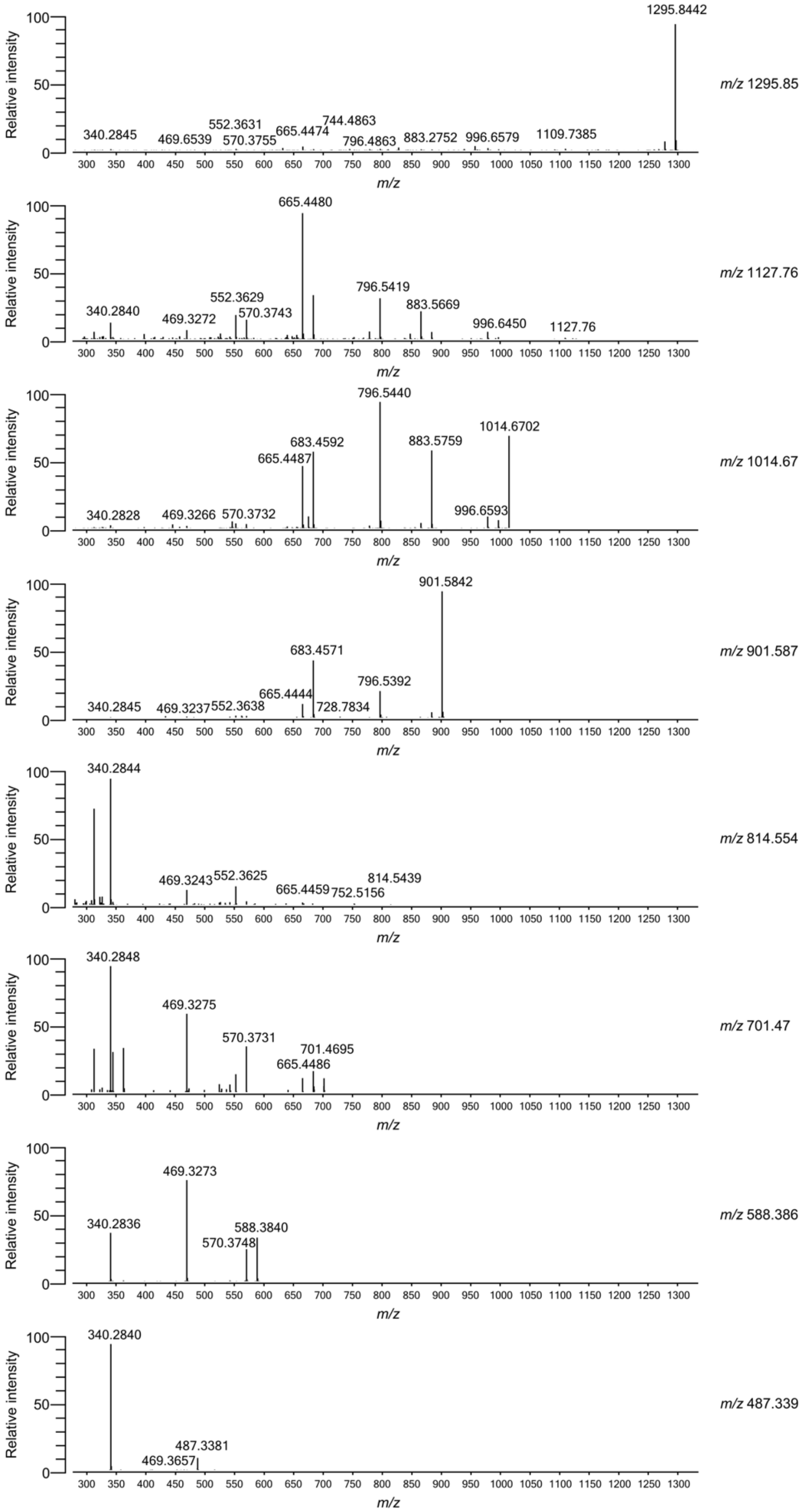


Figure S7 | **Tandem mass spectrometry revealed the relatedness of degradation products to orfamide A.** Observed fragmentation patterns of orfamide A (*m/z* 1295.85) and the degradation products. Similarities between the patterns confirmed the relatedness of each degradation product to orfamide A, representing the loss of amino acids from the C-terminal end of the linearized lipopeptide.
